# Integrative Structural Modeling of Intrinsically Disordered Regions in a Human HDAC2 Chromatin Remodeling Complex

**DOI:** 10.1101/2025.08.08.669391

**Authors:** Jules Nde, Cassandra G. Kempf, Rosalyn C. Zimmermann, Joseph Cesare, Ying Zhang, Jerry L. Workman, Laurence Florens, Michael P. Washburn

## Abstract

Intrinsically disordered regions (IDRs) and intrinsically disordered proteins (IDPs) play pivotal roles in cellular signaling, molecular recognition, and the regulation of various biological processes. These flexible and conformationally dynamic protein segments are difficult to study using structural analysis methods and computational approaches including AlphaFold. Therefore a critical challenge arises when attempting to understand the structural basis of protein-protein interactions involving IDRs. Here we demonstrate that the poorly characterized C16orf87 protein, which we rename as MHAP1, forms a stable complex with HDAC2 and MIER1. These three proteins all contain IDRs whose structure is unknown. We implemented an integrative approach combining experimental crosslinking data with computational modeling techniques (I-TASSER, HADDOCK, AlphaFold) to probe the IDR-driven assembly of the HDAC1:MIER2:MHAP1 complex and build an integrative structural model of this complex. The C-terminal domain of HDAC2, a poorly characterized IDR, promotes interactions between the ELM2 domain of MIER1 as well as the N- and C-termini of MHAP1. These results contrast with most current literature, including the results from AlphaFold alone that are missing structural information on HDAC C-domain. The approach herein can be generalized to study other complexes, emphasizing the need for integrative approaches in determining the 3D structures of IDR/IDP-driven complexes.

## Introduction

Intrinsically disordered regions (IDRs) and intrinsically disordered proteins (IDPs) represent a unique class of macromolecules that lack a fixed, stable three-dimensional structure under physiological conditions ^1^. Despite their structural plasticity, IDRs and IDPs play critical roles in cellular processes such as signaling, transcription, and cellular assembly ^1–4^. The flexible nature of these regions allows them to engage in dynamic interactions with other biomolecules, often serving as hubs for protein-protein interactions ^3,4^. However, the conformational versatility of IDRs complicates efforts to study these interactions using conventional structural techniques, including X-ray crystallography ^5,6^, nuclear magnetic resonance (NMR) spectroscopy ^7–9^, and computational methods like AlphaFold ^10,11^.

The inherent flexibility of these IDRs poses significant challenges for traditional structural biology methods ^6,8,12–14^, as they often fail to capture the full spectrum of conformational states these proteins can adopt in solution. The difficulty in obtaining high-resolution structural data for IDR-containing complexes underscores the need for advanced integrative approaches ^15^. Techniques that combine experimental data with computational modeling, such as cross-linking mass spectrometry (XL-MS) ^16–19^ coupled with molecular modeling techniques (I-TASSER ^20,21^ and HADDOCK ^22^) or deep learning approaches like AlphaFold ^10,11^, are now emerging as powerful tools for studying the dynamic behavior of IDPs and IDRs. XL-MS combines chemical cross-linking with mass spectrometry to provide valuable structural and functional insights into protein interactions, conformational changes, and the dynamics of protein complexes in their cellular context ^18,23^. Furthermore, XL-MS approaches can be used to build integrative structural models of protein complexes ^16,24,25^.

Examples of proteins with large IDRs include Histone Deacetylase 2 (HDAC2) ^16,17,26–30^, Mesoderm Induction Early Response 1 (MIER1) ^26,28,31^, and Chromosome 16 Open Reading Frame 87 (C16orf87) ^32,33^. HDAC2 is a class I histone deacetylase ^27^ involved in removing acetyl groups from lysine residues of histones, leading to chromatin condensation and transcriptional repression ^29^. This deacetylation process is critical in controlling gene expression and regulating cellular functions such as cell cycle progression, differentiation, and apoptosis ^27,30^. MIER1 is a transcriptional co-repressor that has been shown to play a role in regulating gene expression, often by interacting with other repressive complexes ^26,28,31,34^. MIER1 is a multi-domains protein that includes the ELM2 and the SANT domains, which are both critical for triggering protein-protein interactions ^26^. MIER1 is involved in chromatin remodeling and is a part of multi-protein complexes, where it contributes to transcriptional repression, likely by recruiting other repressive factors such as HDACs through its ELM2-SANT domain ^28^. Independently, MIER1 can function in a variety of biological processes, including early developmental stages ^31,34^. However, its highly disordered nature has complicated the determination of its 3D structure, either experimentally or by using advanced computational approaches. Lastly, C16orf87 is a poorly characterized protein whose function remains unknown but it likely interacts with chromatin remodeling proteins like HDAC2 ^32,33^.

The structures of both MIER1 and C16orf87 remain unsolved experimentally likely due to their large content in IDRs, and as such their 3D structures are currently only predicted with AlphaFold ^10,11^. HDAC2 3D structure shows a region of high confidence in AlphaFold, but a long C-terminal loop beginning with Histidine 376 and ending with the C-terminus (Pro488) is unstructured with nearly all aminos acids positions of low to very low prediction confidence. Despite the impressive advances of AlphaFold in protein structure prediction, its limitations in accurately modeling IDRs are evident ^13,15^. IDRs contribute to the challenge in determining complete and reliable 3D structures of individual proteins as well as protein complexes.

Given these challenges, an integrative approach that combines computational models with experimental data is critical for providing more complete structural insights into proteins and protein complexes. In this study, we first demonstrate that C16orf87 forms a complex with HDAC2 and MIER1. We next used AlphaFold ^10,11^ as a stand-alone approach to predict the 3D structures of the dimeric and trimeric complexes. The results were in line with AlphaFold limitations ^10,11^ in capturing the highly flexible IDRs within the proteins, leading to a low-resolution 3D structure of the complexes. Next, we implemented an integrated approach based on experimental XL-MS data and computational modeling to investigate the 3D structures of individual proteins as well as protein complexes. We revealed not only the individual structures of MIER1 and C16orf87/MHAP1 but also the intricate molecular architecture of the ternary HDAC2:MIER1: MHAP1 complex. Our results showed that the ELM2 domain of MIER1 interacts with HDAC2 through the IDR in the C-domain in agreement with published crosslinking data between HDAC1 C-domain and the ELM2 domain-containing protein RCOR1 ^35^. Integrating experimental techniques, such as cross-linking mass spectrometry, with computational predictions substantially enhances the understanding of the interactions and conformational dynamics of these proteins as well as the role played by the IDRs in triggering protein-protein interactions, particularly in regions that remain unresolved by AlphaFold ^10,11^ alone.

## Results

### Establishing C16orf87 as an HDAC1 and HDAC2 Interacting Protein

Previous Halo-HDAC1 and HDAC2-Halo AP-MS experiments in our lab performed in a HeLa cell line identified C16orf87 as a novel and uncharacterized interactor of HDAC1/2 ^33^. Another study also identified that GFP-HDAC2 interacted with C16orf87 in T-Cells ^32^. To study the interaction of C16orf87 with HDAC1 and HDAC2, we first visualized C16orf87 within the cell using an N-terminally Halo-tagged construct and found that Halo-C16orf87 localized to the nucleus (Suppl. Fig. S1A). We next performed co-localization studies within Flp-In-293 cells transiently co-transfected with either C-terminally Halo-tagged HDAC1 or HDAC2 proteins and C16orf87 with an N-terminal SNAP-Flag (SNAP-F) tag (Fig. 1A). Both HDAC1/2 and C16orf87 proteins co-localize to the nucleus and their localization appears to overlap in cells where both constructs were expressed.

**Figure 1.**
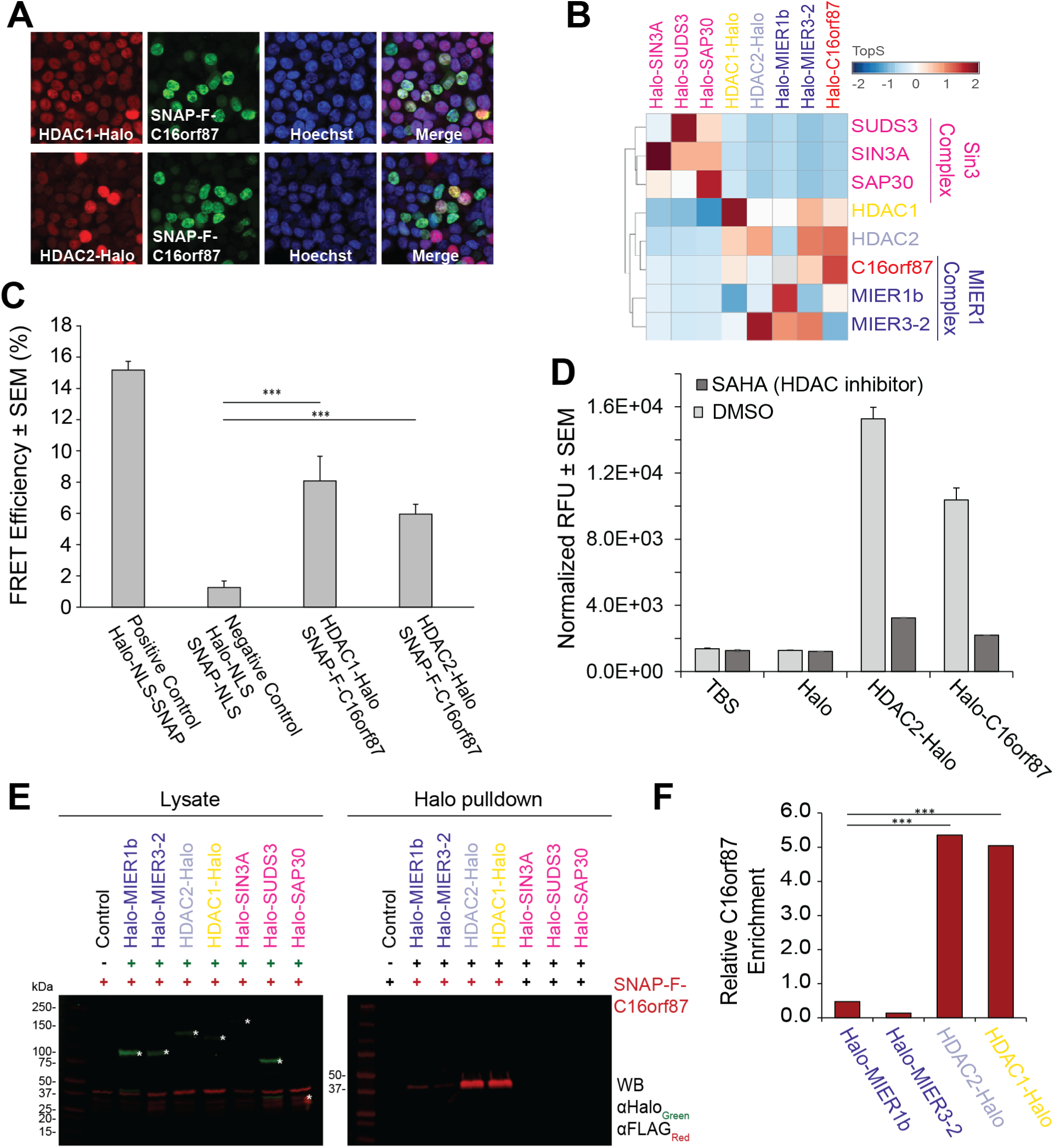
C16orf87 interacts with HDAC1/2 and members of the MIER complex. **A.** Nuclear localization of SNAP-Flag-C16orf87 in Flp-In^TM^-293 cells stably expressing HDAC1-Halo or HDAC2-Halo. Halo-tagged proteins are labeled with HaloTag® TMR ligand (red); SNAP-tagged proteins are labeled with SNAP-Cell ® 505-Star ligand (green); nuclei are stained with Hoechst dye (blue). Nuclear localization was also observed with a Halo-tagged C16orf87 construct (Suppl. Fig. S1A). **B.** Bait-specific hierarchical clustering on Topological scores (TopS) including the 8 Halo-tagged baits: C16orf87, MIER1b, MIER1-2, HDAC1, HDAC2, SIN3A, SUDS3, and SAP30. Red corresponds to high TopS values and blue corresponds to negative TopS values (Suppl. Table S1B). **C.** AP-FRET analysis between SNAP-F-C16orf87 (labeled with SNAP-Cell® 505-Star ligand) and HDAC1/2-Halo (labeled with HaloTag® MR ligand). The positive control was a Halo-NLS-SNAP construct to measure the FRET efficiency of Halo and SNAP when close together; the negative control confirmed an absence of FRET in cells expressing NLS-Halo and NLS-SNAP on their own. Unpaired t-tests were used for statistical analyses where ***: p≤ 0.0001. Representative images of acceptor and donor fluorescence are provided in Suppl. Fig. S1D, while all measurements and Image J processing are reported in Suppl. Table S2A. **D.** Deacetylase activity assays were performed using equal volumes of eluates from the Halo-only, HDAC2-Halo, and Halo-C16orf87 affinity purifications. Deacetylase activity was measured by a colorimetric reaction, and relative fluorescence unit (RFU) values were normalized to total protein content in each sample as measured by BCA protein assay. Assays were performed in triplicate with error bars representing SEM. Experimental workflow, acquisition setting, raw measurements, and calculations are reported in Supp. Table S2B. **E.** HEK293T cell lysates expressing tagged C16orf87 with or without 7 Halo-tagged baits (MIER1b, MIER1-2, HDAC1, HDAC2, SIN3A, SUDS3, and SAP30) were used for Halo pulldown analysis. Equal volumes of eluate were analyzed by SDS PAGE and Western blotting. SNAP-FLAG-C16orf87 was detected using anti-FLAG mouse monoclonal primary antibody and IRD 680 LT labeled goat anti-Mouse secondary antibody. Halo-tagged baits were detected using anti-Halo rabbit polyclonal primary antibody and IRD 800CW labeled goat anti-Rabbit secondary antibody. **F.** Quantification of relative C16orf87 enrichment in HDAC1/2 and MIER protein pulldowns. Normalized values account for differences in C16orf87 concentration between Halo-tagged samples determined by quantitative Western blotting (Suppl. Table S2C). Band intensities were quantitated using Image Studio v5.2 (Li-Cor). Unpaired t-tests were performed for statistical analyses where *** = p≤0.0001.

Next, an AP-MS analysis of the affinity-purified proteins from HEK293T cells transiently expressing Halo-tagged C16orf87 identified several MIER complex members in addition to other HDAC1 and HDAC interacting proteins (Suppl. Fig.S1B, Suppl. Table S1A). To determine the potential C16orf87 relationship with the MIER complex, we performed reciprocal AP-MS experiments using members of the MIER complex (MIER1b and MIER3-2) (Suppl. Fig. S1B, Suppl. Table S1A). We also compared the MIER and C16orf87 co-purified proteins with previous results obtained using members of the SIN3 complex (SAP30, SIN3A, and SUDS3), as well as HDAC1-Halo and HDAC2-Halo as baits (Suppl. Fig. S1B, Suppl. Table S1A). The selection of these baits allowed us to assess the specificity of C16orf87 interaction with the MIER complex. We performed a Cystoscape analysis, in which we identified protein interactions that are significantly enriched (log_2_FC >2, FDR_up_ <0.05) with at least one of the bait proteins. In this analysis, we identified a significant enrichment of C16orf87 to the MIER complex members (Suppl. Fig. S1C). Additionally, the MIER complex members (MIER1b, MIER3-2) reciprocated this close relationship (Suppl. Fig. S1C). This result was further confirmed with hierarchical clustering, in which C16orf87 distinctly clusters with MIER complex members, while the three SIN3 complex members did not (Suppl. Fig. S1B). Lastly, in addition to the traditional clustering analysis, we also calculated Topological Scores (TopS) ^36^ across these AP-MS analyses (Suppl. Table S1B). This complementary scoring method can further cluster proteomics data, in which a high TopS value suggests direct protein-protein interactions (Fig. 1B). We found that, similarly to hierarchical clustering data (Suppl. Fig. S1C), TopS analysis (Fig. 1B) identified a close interaction relationship between C16orf87 and MIER complex members that is distinct from that of the SIN3 complex members.

To further validate the interaction between C16orf87 and HDAC1 and HDAC2, we next implemented acceptor photobleaching fluorescence resonance energy transfer (AP-FRET), an *in vivo* imaging approach. We used the donor/acceptor pairs of SNAP-F-C16orf87/HDAC1-Halo and SNAP-F-C16orf87/HDAC2-Halo co-transiently transfected into HEK293T cells (Suppl. Fig. S1D, Suppl. Table S2A). We detected the pairwise interaction between C16orf87 and HDAC1/2 by determining the signal increase following photobleaching of HDAC1-Halo or HDAC2-Halo in the acceptor cells (Fig. 1C). Our results showed a significant increase in FRET efficiency compared to negative control, demonstrating that the two SNAP-F-tagged and Halo-tagged proteins must have been within <10 nm of each other (Fig. 1C). This experiment confirms that both HDAC1 and 2 directly interact with C16orf87 *in vivo*.

To determine whether the HDAC1 and 2 associated with C16orf87 were enzymatically active, we performed a fluorogenic histone deacetylase activity assay. This assay functions in two enzymatic steps (Suppl. Table S2B). The first step consists of the deacetylation of ε-acetylated lysyl moieties in the presence of HDAC activity. If deacetylation occurs, then a second trypsin cleavage step releases a measurable fluorescent compound ^17,37^. We purified the proteins co-eluting with HDAC2-Halo and Halo-C16orf87 and implemented this HDAC activity assay with the HDAC inhibitor SAHA serving as a control. We found that C16orf87 did co-purify with an enzymatically competent HDAC2 (Fig. 1D, Suppl. Table S2B). This finding suggests that C16orf87 interacts with an active HDAC1/2-complex, as deacetylase activity is maintained.

To further validate the direct interaction of C16orf87 with the MIER complex, we performed a series of pulldowns on HEK293T cells co-transiently expressing SNAP-F-tagged C16orf87 and Halo-tagged MIER1b, MIER3-2, HDAC1, HDAC2, SIN3A, SAP30, and SUDS3 (Fig. 1E). The results of these Halo-immunoprecipitations confirmed the association of C16orf87 with HDAC1, HDAC2, MIER1b, and MIER3-2 (Fig. 1E). However, measuring the intensity of the C16orf87 bands by quantitative Western blotting (Suppl. Table S2C) clearly revealed a greater enrichment of C16orf87 with both HDAC1 and 2 compared to the 2 MIER complex members (Fig. 1F). On the other hand, no C16orf87 association with the SIN3 complex members was observed, as predicted by the AP-MS results (Fig. 1B, Suppl. Fig. S1B-C). This confirms C16orf87 as a *bona fide* member the MIER complex, with a tight association to HDAC1 and HDAC2. Based on the relationships characterized through biological validation of C16orf87 so far, we termed C16orf87 as **<u>M</u>**IER1-**<u>H</u>**DAC **<u>A</u>**ssociated **<u>P</u>**rotein **<u>1</u>**, or MHAP1. From this point forward we will refer to C16orf87 as MHAP1.

### AlphaFold-Generated Combinatorial Assemblies of HDAC2, MIER1, and MHAP1

We next sought to determine how the HDAC2:MIER1:MHAP1 complex might assemble. Using a combination of AlphaFold3 (AF3) ^10^ —which considers the pairwise interactions between subunits of a complex for improved 3D modeling of protein complexes— and AlphaFold-Multimer (AFM) ^38^ —a version of AlphaFold specifically designed for predicting protein complexes— we built structural models of the three possible dimers (Fig. 2A). We also modeled the structure of an HDAC2:MIER1:MHAP1 trimeric complex, into which one copy of each protein is assembled (Fig. 2B). We only performed the comparative analyses described below on the top scoring models for the 3 dimers and the trimer since good structural alignment was observed for all dimers when superimposing the top five scoring models predicted for each dimer (Suppl. Fig. S2A). The slight variations between these top scoring models suggest that the complexes may exhibit conformational adaptability, potentially allowing for functional changes in response to cellular signals or additional molecular interactions.

**Figure 2.**
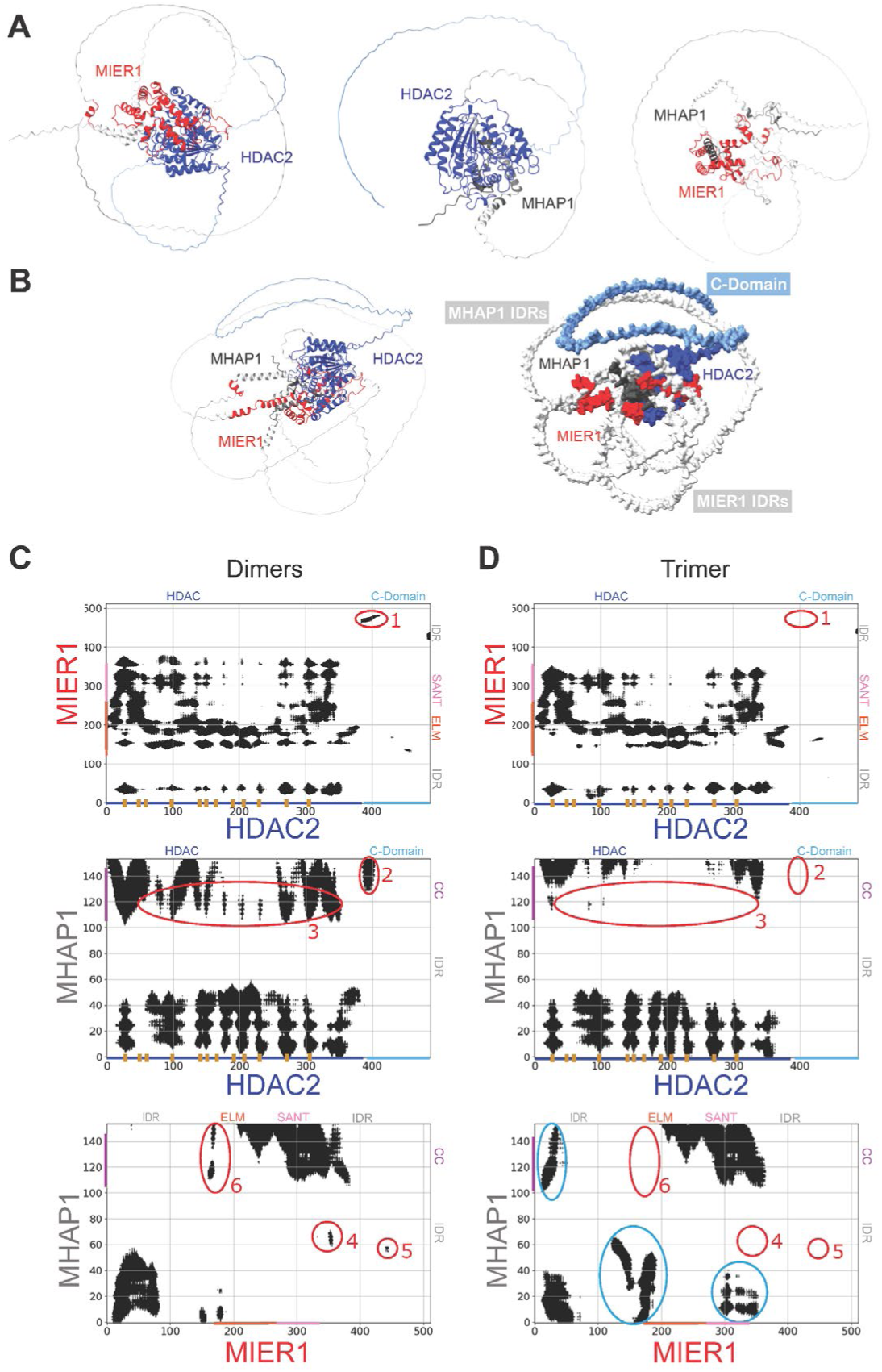
AlphaFold-Based Dimeric and Trimeric Models of Complexes Containing HDAC2, MIER1, and MHAP1. **A.** Structural models of the MIER1:HDAC2, MHAP1:HDAC2, and MHAP1:MIER1 dimers predicted by AlphaFold. Comparisons of the top five assemblies predicted for each dimer are also provided in Supp. Fig. S2A. **B.** Structural model of the trimeric complex as predicted by AlphaFold, with density representation of the complex on the right. MIER1 and MHAP1 IDRs are in light gray, HDAC2 C-domain in light blue, and the ligand binding sites within the dark blue HDAC-domain in yellow. **C-D.** Residue contact maps highlighting critical hotspots for each binary interaction in each dimer and within the trimer. Regions circled and numbered in red denote differences between binary interactions within the heterodi-and tri-meric complexes, while areas circled in blue are unique to the trimer. The corresponding heatmaps of lysine interactions within AF dimers and trimer are plotted in Supp. Fig. S2B.

To simplify the comparison of the 3D structural models between the dimeric and trimeric complexes, we plotted 2-dimensional contact maps of residues within 30 Å of each other in the dimers (Fig. 2C) and trimer (Fig. 2D), as well as heatmaps of the Euclidean distances measured between the α-carbons of all lysine residues (Suppl. Fig. S2B). Overall, the contact maps and interaction analysis between lysine residues revealed several key contacts between known structural domains and IDRs and emphasized the clustered nature of interactions between any two subunits, suggesting highly selective binding interfaces.

In the HDAC2:MIER1 dimer, most of the contacts within this defined distance threshold were observed between the N-terminal HDAC domain and the ELM2-SANT domain, in agreement with previous studies that have shown that MIER1 and HDAC1 interact with each other through their ELM2 and residues 1-376, respectively ^28^. These contacts were reproducibly maintained in the trimeric assembly (Fig. 2D, Suppl. Fig. S2B). The spatial proximity of interacting residues between HDAC2 and MIER1 emphasizes the functional relevance of these domains that are likely to contribute to the specificity and affinity of the binding interface.

In the HDAC2:MHAP1 dimer, the N-terminus, IDR, and C-terminal coiled-coil (CC) regions of MHAP1 interacted extensively with the N-terminal HDAC domain of HDAC2 (Fig. 2C, Suppl. Fig. S2B), potentially playing a critical role in stabilizing the interface of the complex. In the trimer model, the interaction between MHAP1 N-terminus and the HDAC domain was maintained, however, the range of contact between MHAP1 CC domain and the HDAC domain was significantly reduced (areas circled in red and numbered as “3” in Fig. 2D, Suppl. Fig. 2B), hinting at structural rearrangements in the trimer.

In the MIER1:MHAP1 dimer, the network of interactions between residues was concentrated in specific regions, with MHAP1 N-terminus and CC domain interacting with MIER1 N-terminal IDR and ELM2-SANT ^26^ domains, respectively (Fig. 2C, Suppl, Fig. 2B), suggesting that these areas are again crucial for maintaining the structural integrity and functionality of the dimer. These contacts were indeed conserved in the trimeric complex (Fig. 2D). Therefore, MIER1 might interact with MHAP1 through its highly conserved ELM2-SANT domains.

While the N-domain of HDACs promotes their deacetylase activities, their unstructured C-domain (the last 100+ residues) has been shown to also promote protein-protein interactions ^27–30,35^. However, since the C-domain is missing or left “unmodeled” in all HDAC1/2 experimentally resolved structures, AlphaFold tends to recognize the N-domain of HDACs (residues 1-376) as the only region where protein-protein interactions might occur. Here, we observed a small proximal section of HDAC C-Domain, containing lysine 404, in proximity to the very C-termini of MIER1 and MHAP1 in their respective dimer models (labeled as “1” and “2” in Fig. 2C and Suppl Fig. 2B). Additional isolated contact points were measured between MIER1 N- and C-terminal IDRs and MHAP1 IDR and CC regions (labeled as 4-6 in Fig. 2C and Suppl Fig. 2B). However, none of these interactions were observed when the binary contact maps and heatmaps of lysine distance were plotted for the AF trimer model (Fig. 2D, Suppl. Fig. 2B), indicating that they were spurious and likely due to the flexibility of the long unstructured regions in the AF models.

Strikingly in the trimeric model, extensive regions of contact were newly observed between MHAP1 N-terminal IDR and MIER1 N-terminal IDR, ELM2 and SANT domains, as well as MHAP1 CC domain and MIER1 N-terminal IDR (contact densities circled in blue in Fig. 2D, Suppl. Fig. 2B). The contact map between MIER1 and MHAP1 showed critical hotspots that bridge the proteins together within the trimeric assembly, specifically through the SANT domain of MIER1 and the N- and C-termini of MHAP1. The contact map between HDAC2 and MHAP1 revealed complementary interactions that help anchor the third protein within the trimer through the interactions between the HDAC2 residues 1-376 and the N- and C-termini of MHAP1. These contact maps showcase that specific domain interactions between all three proteins might contribute synergistically to the formation of the complex heterotrimeric complex. The surface representation highlights the interface regions and the overall molecular shape of the trimer (Fig. 2B), with HDAC2 and MIER1 forming the core of the complex and MHAP1 primarily contributing to structural stability.

Furthermore, the presence of HDAC2 in the trimeric complex led to a notable conformational change in MIER1, which adopted a more compact structure upon its interaction with HDAC2. This transition is evident in the change in MIER1 when compared to its individual model, indicating a folding mechanism commonly seen in intrinsically disordered proteins ^2–4^. This "binding and folding" or induced-fit phenomenon ^1,39^ suggests that MIER1 undergoes structural adjustments upon binding, a hallmark of IDPs, allowing it to engage more effectively in the complex. Additionally, MHAP1, due to its small size and flexible nature due to the presence of IDR and coil-coiled domains, plays a structural role in the trimeric complex. Its involvement appears to stabilize the interaction between HDAC2 and MIER1, while its inherent flexibility allows the complex to undergo conformational changes. This adaptability is essential for the complex potential to readjust or adapt to new cellular conditions, facilitating its dynamic response to varying biochemical environments ^3^.

Overall, while AlphaFold, and its improved versions, AF3 and AFM ^10,11,38^, was able to assemble a putative structure for the HDAC2:MIER1:MHAP1 heterotrimer, large areas of each protein remain unstructured and disordered (Fig. 2B).

### Integrative Structural Model of the Trimeric HDAC2:MIER1:MHAP1 Complex

Since the structures of MIER1 and MHAP1 have not been resolved experimentally, AlphaFold was unable to predict high-resolution structures for these proteins due to their intrinsic flexibility and high IDR content. Therefore, we implemented an integrative structural modeling (ISM) approach (Suppl. Fig. S3A) that combines multiple data sources, modeling methods, and experimental data to predict the structure of proteins or protein complexes ^16,40–42^ ^16,24,25^. Such integrative method can capture dynamic and conformational flexibility, which might be missed by multiple sequence alignment approaches like Alpha-Fold ^41^.

First, we performed XL-MS analyses of three biological replicate affinity purifications of N-terminally Halo-tagged MHAP1 transiently expressed in HEK293T cells. Combining the replicates, a total of 525 crosslink spectrum matches (CSMs) were annotated (Suppl. Table S3A), corresponding to 165 and 52 unique intra- and inter-molecular crosslinks, respectively (Suppl. Table S3B), mapped to 148 proteins (Suppl. Table S3C). In this dataset, unique crosslinks between MIER1 and MHAP1 and between HDAC2 and MHAP1 were detected and identified (Suppl. Fig S3B).

Only 4 of the 20 lysine residues in the MHAP1 sequence (Q6PH81:CP087_HUMAN) were found to be crosslinked to other lysine residues. All four of these residues were in the intrinsically disordered and unstructured loop in the AlphaFold prediction of the UPF0547 protein C16orf87/MHAP1 structure and their per-residue model confidence scores (pLDDT) were in the low to very low range (Suppl. Fig. S3C). Next, we utilized experimental information from the XLMS data to guide the computational prediction of each protein. The XL-MS data provided valuable spatial constraints that were incorporated into individual models computed by i-TASSER ^20^ (Suppl. Fig. S3A), which resulted in significantly improving predicted structures especially within the IDRs, now strikingly predicted to fold into α-helices for all three proteins (light gray and light blue regions in Fig. 3A).

**Figure 3.**
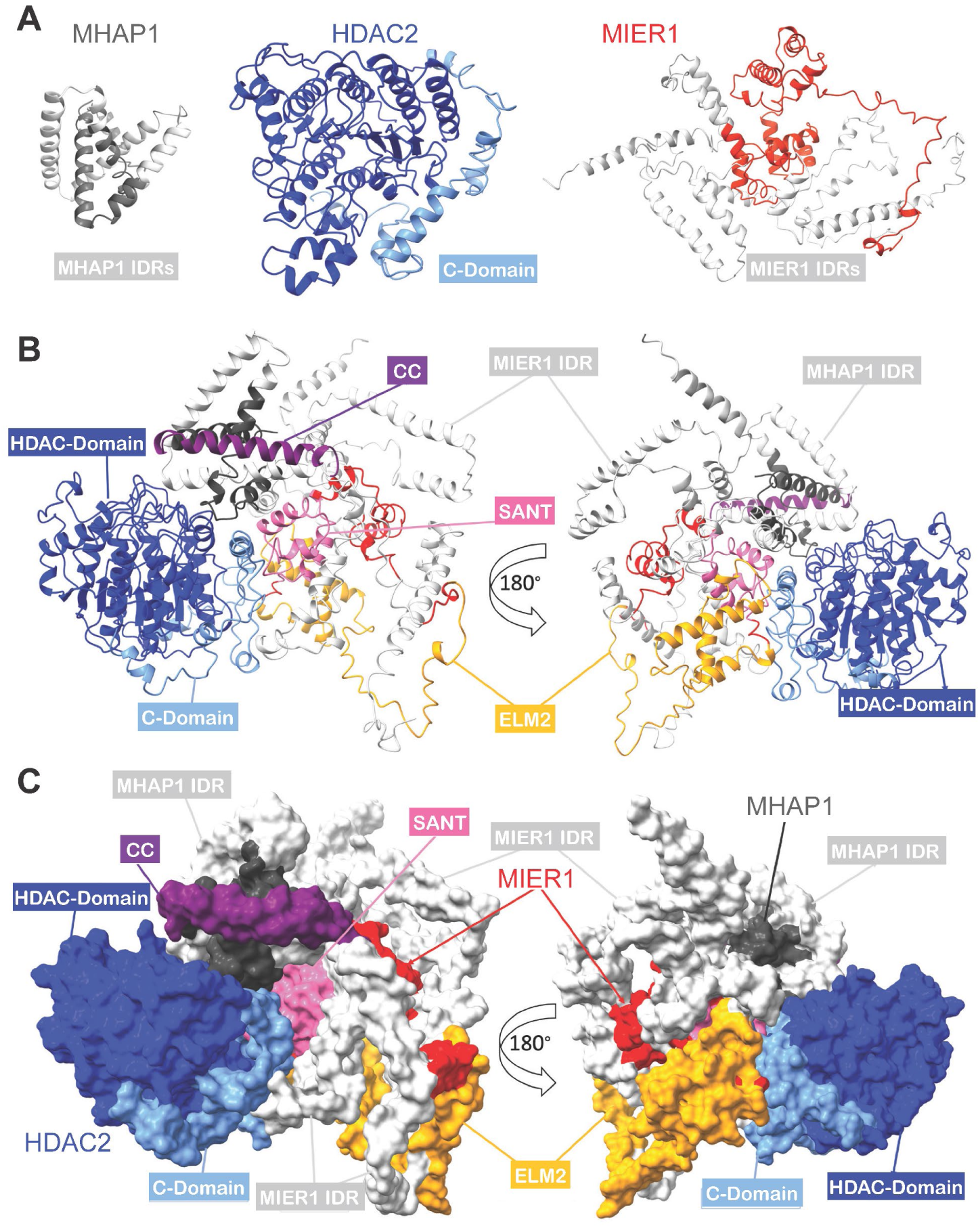
Integrative Structural Model of the HDAC2:MIER1:MHAP1 Trimer. **A.** iTASSER predicted models (after workflow step IV in Fig. S3A) for HDAC1, MIER1, and MHAP1 (visualized in Chimera as ribbon representation) used for assembling the heterotrimeric complex in steps V and VI in Fig. S3A. **B-C**. Ribbon and space-filled formats of ISM trimer with 180° rotation on the right. In all panels, each of the complex subunits and their domains and regions is color-coded: HDAC2 in blue, MIER1 in red, MHAP1 in dark gray; HDAC2 C-terminal domain in light blue; MIER1 ELM2 and SANT domains in orange and pink, respectively; MIER1 and MHAP1 IDRs in light gray; MHAP1 coiled-coil (CC) region in purple (see Supp. Video 2).

After integrating experimental distance constraints derived from intra-molecular crosslinks for the computational modeling of the individual subunits, we used CPORT ^43^ in combination with HADDOCK ^22^ to predict their docking interfaces and assemble the HDAC2:MIER1:MHAP1 trimer (Fig. 3B-C, Suppl Fig. S3D). In contrast to the AF model (Fig. 2B), we observed a structural shift as both MIER 1 and MHAP1 interact with HDAC2. More specifically, MIER1, through its ELM2 domain, and MHAP1, through its N- and C-termini, interact with HDAC2 C-domain (Fig. 3B-C). This is agreement with the C-domain promoting protein-protein interaction, as previously shown by crosslinking and electron microscopy structural analysis of HDAC1 in a ternary complex with LSD1 and RCOR1^35^. These observations underscore the importance of XL-MS data in guiding 3D structural prediction.

Again, we plotted contact maps of residues within 30 Å of each other between any two subunits of the complex (Suppl. Fig. S4A), as well as the heatmap of all Euclidian distances between lysine residues (Suppl. Fig. S4B), to better visualize interaction interfaces at the molecular level. The lysine residues form well-isolated hotspots from all three proteins that are essential for the overall stability and the structural integrity of the complex (Suppl. Fig. S4B). The contact maps between MIER1 and MHAP1 reveal important domain interactions (Suppl. Fig. S4B, circled in blue) that help bridge these two proteins together, through MIER1 SANT domain and the N- and C-termini of MHAP1, contributing to the overall stability of the complex. The interactions between HDAC2 and MHAP1 are highlighted through key hotspots (Suppl. Fig. S4B, circled in blue), which demonstrate how the smaller MHAP1 helps anchor HDAC2 in the complex (Suppl. Fig. S4C). To better visualize these specific interaction interfaces, each named domain was color-coded, while IDRs were left in light gray (Fig. 3B-C). Interactions between HDAC2 and MIER1 occur through key contacts between MIER1 ELM2-SANT (Fig. 3B-C, orange and pink, respectively) and HDAC2 C-domain (Fig. 3B-C, light blue).

A key aspect of this ISM model is the IDRs forming helical secondary structures (Suppl. Table S4). In MIER1, the three IDRs are folding into multiple distinct helices, and the same is true for MHAP1 internal intrinsically disordered loop. These IDRs are likely a composition of short linear motifs (SLMs) and intrinsically disordered domains (IDDs). The large and poorly characterized HDAC2 C-terminal domain is around 100 amino acids long and is likely a molecular recognition feature (MRF) domain. Strikingly, in the ISM model trimer, the C-domain is now folded into six short α-helices that form a compact structure (Fig. 3B-C, Suppl. Table S4). The final 3D structure of the HDAC2:MIER1:MHAP1 complex (Fig. 3B-C) shows direct contacts between the no-longer disordered C-domain (light blue) and MIER1 ELM2 domain (orange). Furthermore, the C-domain folds back to promote protein-protein interactions, not only between HDAC2 and MIER1, but also between HDAC2 and MHAP1, which illustrates the folding-upon-binding mechanism, first reported for pKID, a disordered region within the transcriptional co-activator CREB ^44^ and favored to explain IDRs ability to form protein interactions with a variety of partners.

Overall, despite challenges in predicting high-resolution structures for MIER1 and MHAP1 individually, our integrative approach has led to a model of the entire trimeric complex where each protein is completely modeled into a globular structure (Suppl. Fig. S3D). Our integrated 3D model reveals how the three subunits form a stable and compact trimer, with HDAC2 and MIER1 forming the core (Suppl. Fig. S4C), and MHAP1 playing a supporting role in stabilizing the complex. The overall architecture of the complex highlights the structural coordination between the proteins.

### Comparison of the 3D Models Generated by AlphaFold and Integrative Structural Modeling

Based on the HDAC2:MIER1:MHAP1 trimers described in the previous two sections, the AF and ISM models do not agree on the interacting interfaces within these assemblies. To better assess these structural differences, we superimposed the AF and ISM trimeric models using the HDAC domain, the one section with a resolved crystallographic structure, as anchor (Fig. 4A and Supp. Videos 1 and 2). The structural differences are significant and fall into two main categories: the quaternary organization of the subunits within the complex and the secondary and tertiary structures of each protein.

**Figure 4.**
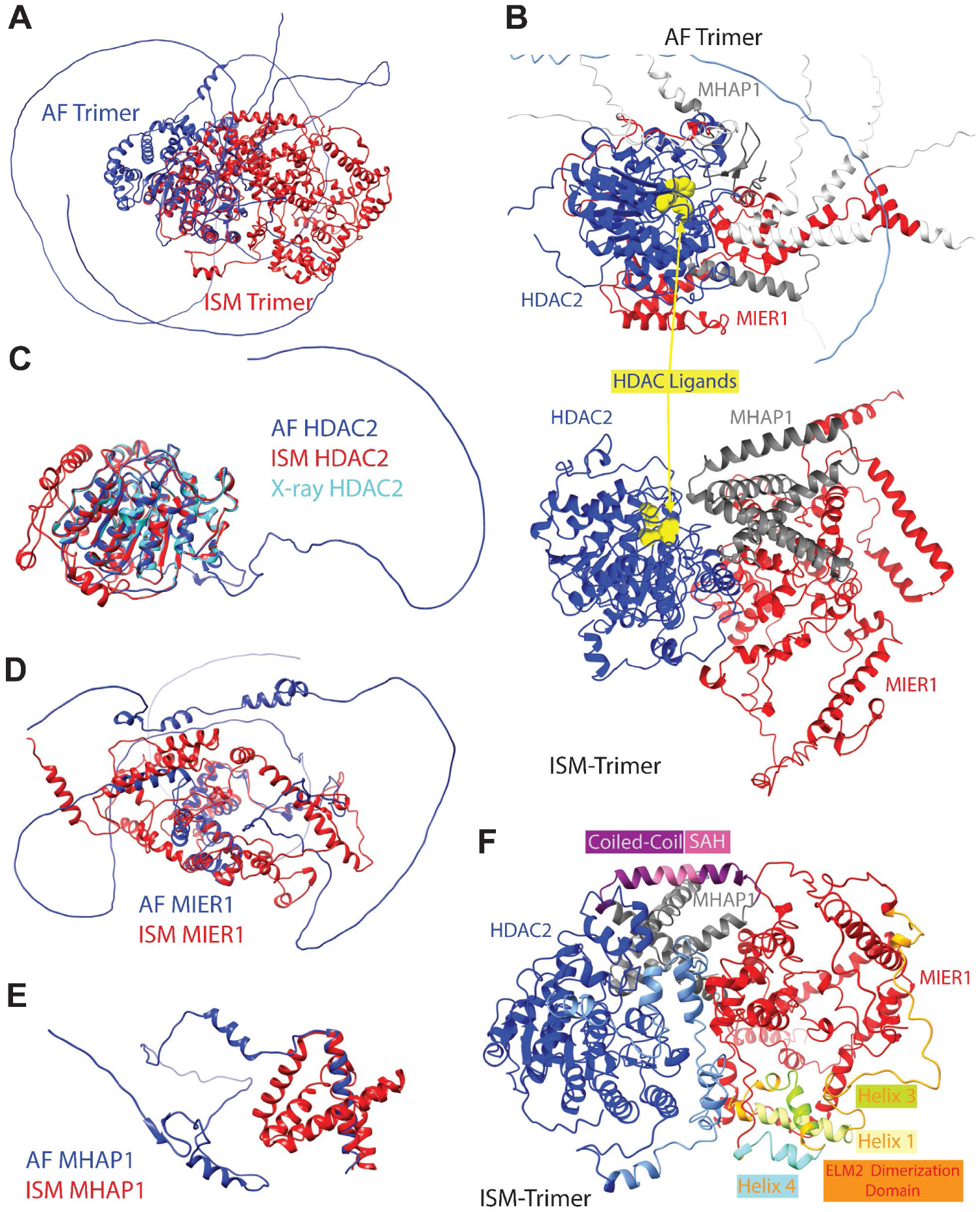
Comparative Analysis of the AlphaFold-Based and the Integrative Structural Models. **A.** The 3D structures obtained through integrative structural modeling (ISM, red) and AlphaFold (AF, blue) for the HDAC2:MIER1:MHAP1 trimeric complex are superimposed using the HDAC domain of HDAC2 as anchor. **B.** Location of ligands, in yellow, within the HDAC domain in the AF and ISM models. **C.** HDAC2 3D structures from AF (blue), ISM (red), and 7ZZ0.pdb (cyan) were superimposed aligning their respective HDAC domains. **D.** AF and ISM models of MIER1, illustrating AF limitations in predicting unstructured regions or regions lacking experimental data. **E.** Structural comparison of MHAP1 models predicted, in which the long C-terminal helix common to both models was used to align the structures in 3D. **F.** 3D model of the HDAC2:MIER1:MHAP1 complex obtained using ISM, in which the 3 helices within MIER1 ELM2 dimerization domain and MHAP1 Coiled-Coil/Stable Alpha Helix domain are highlighted.

First, a drastic shift was observed in which of HDAC2 two domains was responsible for interacting with the other two subunits. On the one hand, AF predicted all interactions to involve the catalytic HDAC domain (Fig. 2D and Suppl. Fig. S2B); on the other hand, ISM predicted the tightest interactions at the core of the complex to be through the C-domain (His 376 – Pro 488) (Fig. 3B-C and Suppl. Fig. S4D). These different quaternary architectures influence the accessibility of the substrate binding channel within the HDAC domain. In the AF-predicted trimeric complex, the known ligand binding loops of the HDAC domain are not accessible due to its expansive interactions with MIER1 and MHAP1 (Fig. 4B and Suppl. Fig. S2B). This would entail that HDAC is not enzymatically active in this complex, which disagrees with our experimental validation (Fig. 1D) that MHAP1 pulls down an active HDAC1/2 complex. In our ISM model, MIER1 has very limited contact points with the HDAC domain (Suppl. Fig S4A), which allows for the ligand binding sites to be fully accessible in the quaternary assembly (Fig. 4B). Therefore, the subunit organization within the ISM model is in biological agreement with a fully functional HDAC2:MIER1:MHAP1 complex competent in histone deacetylation.

Second, the IDRs in each of the proteins transitioned from large intrinsically disordered tails and loops “flopping around” without any contact with the core of the complex into canonically folded secondary structures (α-helices) involved in tertiary interactions (“folding-back”) with the globular core of the complex (Fig. 4A and Supp. Videos 1 and 2). Zooming in to the individual subunits, we compared the 3D models predicted by AlphaFold and ISM against one of the experimentally resolved HDAC2 structures available in the protein databank (7zzo.pdb) ^45^. The two models showed good agreement with the experimentally determined structure of the HDAC domain (Fig. 4C), with near perfect alignment of the α/β fold with 13 α-helices surrounding a central 8-strand parallel β-sheet ^46^. This again reflects AF ability to align ordered domains or structures with known experimental data. However, AF modeled a long loop with no secondary structure in the unstructured HDAC2 C-domain, further emphasizing its challenges in predicting flexible or disordered regions. The integrative approach, incorporating experimental distance constraints, provides a refined structure for the C-domain, which now contains six α-helices from residues 389-486 (Suppl. Table S4). Furthermore, in the ISM model, the C-domain is folded back onto the α/β fold of the HDAC domain, hence providing an interaction interface for the other two proteins that was absent from both the AF trimer and the experimentally solved HDAC2 structure. MIER1, being an intrinsically disordered protein, presented a challenge for AlphaFold. In addition to N-terminal and long C-terminal unstructured tails and an internal IDR loop, AF predicted several short helical turns and helices between the first 2 IDRs (Fig. 4D and Suppl. Table S4). AF predicted a series of mostly helical regions within the ELM2 domain starting with a long string of misshapen helix/helical turns followed by 3 short helices (Fig. 4D and Table S4). The MIER1 SANT domain is likely to adopt the typical three-helix bundle structure at the core of this domain, which are arranged in a helix-turn-helix motif, while a small hydrophobic core stabilizes this three-helix bundle structure ^47^. Indeed, AF modeled a bundle of three α-helices within MIER1 SANT domain, likely because multiple experimental reference structures are available for SANT domains ^48^. The AF model for MIER1 is not compact with most of the predicted secondary structures separated from each other, except for the SANT three-helix bundle, which served as anchor for the superimposition (Fig. 4D). In sharp contrast, the ISM MIER1 model is more compact, without any of the extended disordered tails and loops (Fig. 4D and Table S4). The IDR in MIER1 between amino acids 73 and 175 now is formed into two primary α-helices (Fig. 4D and Table S4). In addition, MIER1 IDR between amino acids 366-512 now is formed into 6 α-helices prior to the C-terminal tail (Fig. 4D and Table S4).

As with HDAC2, AlphaFold performed reasonably well in predicting MHAP1 ordered regions, modeling a short N-terminal helical turn (AAs 8-11), followed by a few short β-strands (AAs 12-35), an α-helix (AAs 110-121) in the coiled-coil region, and one long C-terminal α-helix (AAs 130-151) (Table S4). AF again modeled MHAP1 IDR region as a long unstructured loop (AAs 49-109) with a short helical turn from AAs 65-67 (Fig. 4E and Suppl. Table S4). These secondary structures and unstructured loop did not appear to have any interactions but existed as separate entities that failed to fold into a globular structure. (Fig. 4E). The superimposition of the long C-terminal α-helix common to the 2 MHAP1 models (Fig. 4E) showed that the boundaries of the long looping IDR predicted by AF have folded into a short helix and three additional α-helices (Fig. 4E and Table S4). These α-helices further interact with the coiled-coil and C-terminal helices to form a tightly packed, globular bundle (Fig. 3A and 4E), a likely accurate representation of this small protein.

Beyond the regions annotated as intrinsically disordered, our ISM model provides additional insights into the capacity of MIER1 ELM2 domain and MHAP1 coiled-coil region in supporting dimerization. ELM2 domains, such as the one in MIER1, are also known to lack regular secondary structures when expressed on their own, such as a construct expressing the MTA1 ELM2 domain only interacted with HDAC1 when co-transfected ^49^. In the AF model of MIER1 sequence, three α-helices are predicted with varying confidence after the conserved ELM2 motif (Table S4). These align with helices 1, 3, and 4 within MTA1 ELM2 dimerization domain ^49^, while a short 1-turn helix is predicted for AAs 252-252, which could correspond to helix 2 (Table S4). However, in our ISM model of the trimeric assembly, this short helix did not fold further into a longer one (Fig. 4F). As it had been predicted previously for RCOR1 ELM2^49^, which is also lacking this helix 2 due to a short 6 AA stretch between helices 1 and 3, MIER1 ELM2 is also likely to be not competent for dimerization.

Similarly, coiled-coil domains have been involved in oligomerization. In MHAP1, a single 12-residues α-helix was predicted by AF as spanning the coiled-coil domain. CC domains typically comprise multiple helices wound around each other, which is not the case for MHAP1 CC. In our ISM model, this CC-helix was expanded by a couple more turns of helices covering AAs 106-127 (Table S4). Looking into this helix more closely, it has the characteristics of a Single stable α-Helix (SAH) domain defined by uninterrupted helical stretches with specific charged and helical promoting (glutamate, lysine, alanine) amino acid patterns. These amino acid periodicities are used by prediction algorithm such as Waggawagga ^50^, which indeed identified a 14 amino acid SAH window in MHAP1 CC domain (Suppl. Fig. S5B). Unlike coiled-coils, SAH helices are stabilized by intrahelical salt bridges (Suppl. Fig. S5B), which allows them to function as stable single alpha-helices and resist oligomerization. In MHAP1, such SAH helix could then be acting as a rigid molecular spacer (Fig. 3B-C and 4F) to position the functional domains within HDC2 and MIER1 at defined distances.

Our results underscore the importance of incorporating experimental data when predicting the structure of disordered or uncharacterized proteins, where AlphaFold alone may not suffice. While AlphaFold excels in predicting structured domains, we have shown that its predictions for flexible, unstructured intrinsically disordered regions like those C-domain of HDAC2, MIER1, and MHAP1 are significantly less reliable. In contrast, the integrative structural modeling approach, which combines computational predictions with experimental data, leads to more compact and well-folded models by constraining the conformations of disordered regions and improving the overall structural prediction of the complex. These critical observations further confirm the importance of the proposed integrative approach in determining the 3D structures of proteins and protein complexes while considering their native-like environment.

## Discussion

Intrinsically disordered proteins (IDPs) are increasingly recognized as vital to cellular functions such as genome maintenance, signaling, regulation, and complex formation ^1–4^. Unlike globular proteins, IDPs lack a stable three-dimensional structure in isolation ^1,2,3^, existing instead as dynamic ensembles of conformations that may fold in response to environmental cues, post-translational modifications, or interactions with other molecules ^3^. Traditional methods like X-ray crystallography ^6^, NMR ^8^, or cryo-EM ^14^ favor well-ordered proteins, often leading to solving tridimensional structures in which the flexible loops have been removed by genetic engineering to facilitate the formation of crystals or to decrease a protein size to make it compatible with NMR molecular weight limitations. Similarly, since AlphaFold ^10,11^ is machine-learning from the alignments of these solved 3D structures and protein domains, it tends to assume the well-ordered domains are the only possible sites of interaction, mostly ignoring IDRs altogether. While flexibility complicates solving the tridimensional structures of IDPs, their IDRs should not be ignored but rather be a key consideration during the functional and structural characterization of such proteins and the multiprotein complexes into which they assemble.

In this study, we validated C16orf87 as a novel interactor of HDAC2 and MIER1 and, as such, renamed it MHAP1 for MIER1-HDAC Associated Protein 1. Using affinity purification mass spectrometry, *in vivo* imaging, and co-immunoprecipitation, we confirmed the formation of an enzymatically competent ternary assembly, which is notable for the high IDR content of each subunit. The newly named MHAP1 contains an extensive disordered region from AAs 43-119 followed by a coiled-coil domain (AAs 104-132). MIER1 is a transcriptional regulator that contains several IDRs, which are thought to facilitate its interaction with a wide range of proteins, including histone deacetylases like HDAC2 ^26^. Many tridimensional structures of HDAC1 and 2 have been reported as part of various complexes. Forty 3D structures have been solved for HDAC2 alone, the first one of a human HDAC2 complexed with an N-(2-aminophenyl)benzamide (3MAX) solved by X-ray in 2010 ^51^ and the last one of HDAC2-CoREST in complex with KBTBD4R313PRR mutant (9DTQ) solved by EM in 2025 ^52^. However, none of these 40 HDAC2 structure contains the last 100 amino acids defining HDAC1/2 intrinsically disordered C-Domain, which was either not expressed to solve the structure or left “unmodeled”.

Given the limitations of traditional techniques in capturing the dynamic nature of IDPs, we implemented an integrative structural approach that combines multiple experimental and computational methods ^12,40,41^ to characterize the newly defined HDAC2:MIER1:MHAP1 complex. Crosslinking mass spectrometry (XLMS) allowed us to define key interaction sites by providing distance restraints across flexible regions^16,18,19,23–25^. These constraints guided computational modeling using tools such as I-TASSER and HADDOCK and enabled us to generate structural models that account for the disordered and dynamic nature of the complex.

We observed that the IDRs of all three proteins in our ISM models were folded into multiple canonically/regularly shaped α-helices. These secondary structures were expanded from the short proto-helical structures predicted by AlphaFold in both monomers and the trimeric assembly. These short helices have been described previously in IDPs as local residual helical structures ^53^ or pre-structured motifs (PreSMos), which are transient secondary structures that may act as binding hotspots ^54^. Such helix-forming molecular recognition features ^55,56^ confer specific conformational propensities ^57^ to these short amino acid sequences that contribute to the folding-upon-binding mechanism (reviewed in: ^58^). Therefore, the AF trimeric model we generated captures the transient conformational state of each protein with such short irregular helices identified within their IDP loops and tails, while the crosslink-guided ISM models for each protein provide as representation of the fully folded 3D structures.

Our findings further showcase how IDRs contribute to both the stability and adaptability of the HDAC2:MIER1:MHAP1 complex. MIER1 likely serves as a scaffold, recruiting HDAC2 and MHAP1 via specific and well-defined domains, while MHAP1, due to its small size and flexibility, may support complex stability and facilitate conformational shifts. Strikingly, HDAC2 C-domain, which represents almost a quarter of its primary sequence yet had never been solved before, is a key element of the trimeric assembly: its helical secondary structures and its tertiary structure folding-back onto the core of the complex provide the main interaction interface guiding the overall quaternary architecture of the HDAC2:MIER1:MHAP1 complex.

Overall, this work illustrates how integrative structural biology—combining experimental techniques like XL-MS with advanced computational modeling—can reveal the architecture and dynamics of IDP-rich protein complexes. Our solving of the HDAC2:MIER1:MHAP1 complex serves as a proof-of-concept for understanding how disordered regions drive functional assemblies that are otherwise inaccessible through traditional structural methods. As our knowledge of IDPs deepens, such integrative strategies will be essential for decoding their roles in cellular regulation and for guiding therapeutic efforts targeting these dynamic systems.

## STAR METHODS

### Materials

Magne^®^ HaloTag^®^ magnetic affinity beads (G7281), HaloTag^®^ TMR fluorescent ligand (G2991), Rabbit anti-HaloTag^®^ polyclonal antibody (G9281), Sequencing Grade Modified Trypsin (V5111), Mammalian Lysis Buffer (G938A), FuGENE^®^ HD Transfection Reagent (E2311), and Protease Inhibitor Cocktail (G6521) were purchased from Promega. Clone FHC02563 containing the HDAC1 open reading frame was purchased from the Kazusa DNA research institute. HEK293T cells (ATCC1CRL-11268™) were purchased from American Type Culture Collection (ATCC). Flp-In™-293 cells (AHO1112) and AcTEV protease (#12575015) were purchased from Invitrogen. SNAP-Cell^®^ 505-Star ligand was purchased from New England Biolabs (Ipswich, MA). Anti-FLAG^®^ M2 mouse monoclonal (F3165), trypsin from porcine pancreas (T4799), Triton™ X-100 (X100), and Hoechst 33258 (94403) solution were purchased from Sigma-Aldrich. Lipofectamine™ RNAiMAX Transfection Reagent (15338500), Opti-MEM™ Reduced Serum Medium (31985062), Pierce™ BCA Protein Assay Kit (23225), and GlutaMAX™ Supplement (35050061) were purchased from ThermoFisher Scientific. IRDye^®^ 800CW labeled goat anti-Rabbit (926-3211) and IRDye^®^ 680LT labeled goat anti-Mouse (926-68020) secondary antibodies were purchased from LI-COR Biosciences. Salt Active Nuclease (HL-SAN, 70910-202) was purchased from ArcticZymes. Disuccinimidyl sulfoxide (DSSO, 9002863) was purchased from Cayman Chemical. Boc-Lys(Ac)-AMC (A8713) was purchased from ApexBio. Fetal Bovine Serum (FBS, PS-FB1) was purchased from PEAK^®^ Serum. Dulbecco’s Modified Eagle Medium (DMEM, 10-013-CV) was purchased from Corning. Hygromycin B (H-1012-PBS) was purchased from AG Scientific.

### Halo Affinity Purification of Proteins Complexes

For Halo purifications prepared for MudPIT mass spectrometry, extracts were incubated with Magne^®^ HaloTag^®^ beads prepared from either 100 µl Magne™ HaloTag^®^ bead slurry (transiently transfected HEK293T cells) or 200 µl Magne™ HaloTag^®^ bead slurry (Flp-In™-293 cell lines) overnight at 4°C. Beads were washed a minimum of four times with buffer containing: either 10 mM HEPES (pH 7.5), 1.5 mM MgCl_2_, 0.3 M NaCl, 10 mM KCl, and 0.2% Triton X-100 (Flp-In™-293 cell lines), or 50 mM Tris·HCl pH 7.4, 137 mM NaCl, 2.7 mM KCl and 0.05% Nonidet^®^ P40 (transiently transfected cells). Bound proteins were eluted by incubating the beads with AcTEV™ Protease (Invitrogen) overnight at 25°C. Eluates were passed through a Micro Bio-Spin™ chromatography column (Bio-Rad) to remove residual traces of beads.

For Halo purifications prepared for XL-MS, extracts were incubated with Magne^®^ HaloTag^®^ beads prepared from 200 µl Magne™ HaloTag^®^ bead slurry and crosslinked with disuccinimidyl sulfoxide (DSSO, Cayman Chemical Company) MS-cleavable crosslinker as previously described ^16,17^. Briefly, DSSO was added to samples while the proteins were immobilized on beads to a final concentration of 5 mM. Samples were incubated at room temperature for 40 min. Reactions were quenched with the addition of NH4CO3 to a final concentration of 50 mM and samples were incubated an additional 15 min. Bound proteins were eluted by incubating the beads with AcTEV™ Protease (Invitrogen) overnight at 25°C.

### Digestion of Proteins for Mass Spectrometry

Samples were prepared for MudPIT mass spectrometry analysis as previously described ^17^. Briefly, purified proteins were precipitated by incubation with 20% trichloroacetic acid overnight at 4°C. The TCA precipitated proteins were concentrated by centrifugation, washed twice in ice-cold acetone, and residual acetone removed using a vacuum concentrator. Proteins were then resuspended with buffer containing: 100 mM Tris·HCl pH 8.5 and 8 M urea. Disulphide bonds were reduced by adding 0.5 mM tris(2-carboxylethyl)-phosphine hydrochloride (TCEP) and incubating samples at room temperature for 30 minutes. Samples were treated with 10 mM chloroacetamide (CAM) for a further 30 minutes to prevent disulphide bond reformation. Denatured proteins were digested with endoproteinase Lys-C for at least 6 hours. The urea concentration was then reduced to 2M using 100 mM Tris·HCl pH 8.5, CaCl2 was added to a final concentration of 2 mM, and proteins further digested with trypsin overnight. Reactions were stopped by adding formic acid (5% final concentration).

### MudPIT Mass Spectrometry and Data Analysis

Digested samples were loaded onto microcapillary columns packed with three immobilized phases (reversed phase resin; strong cation exchange C18 resin; reversed phase resin). Bound peptides were then eluted from the column using an Agilent 1100 or 1200 series quaternary HPLC pump. Peptides were resolved using 10 multidimensional chromatography steps ^59^ and analyzed using linear ion trap (LTQ) mass spectrometers (Thermo Scientific) in positive ion mode. MudPIT data was analyzed as previously described ^17^. Briefly, a minimum of three biological replicates were acquired for each APMS analysis. As a control, cells were generated that stably express HaloTag^®^ alone, or that were transiently transfected with HaloTag^®^ alone. The resulting .RAW files were converted to .ms2 files using RAWDistiller v. 1.0^60^. ProLuCID v1.3.5^61^ was used to match spectra against a database (Genome Reference Consortium Human Build 38 patch release 13) containing 44519 unique protein sequences, 426 of which were contaminant proteins. The database also included shuffled versions of all sequences for false discovery rate (FDR) estimation (Table S1). Database search was performed with a static modification of +57 Da on cysteine residues, a dynamic modification of +16 Da on methionine residues (oxidation), and a mass tolerance of 800 ppm for precursor and fragment ions.

### Crosslinking Mass Spectrometry and Data Analysis

Digested peptides were analyzed with an Orbitrap Fusion™ Lumos™ (Thermo Fisher) coupled to a Dionex U3000 HPLC using previously described data acquisition settings^16,17^. Briefly, Proteome Discoverer v2.4 and the XlinkX node (Thermo Fisher) ^62^ were used to identify crosslinked peptides from .RAW files. For each .RAW file, XlinkX was used to identify MS2 fragmentation scans with reporter ions characteristic of DSSO crosslinked peptides. Peptides were searched against a human proteome database (Genome Reference Consortium Human Build 38 patch release 13) containing 44519 unique protein sequences, 426 of which were contaminant proteins, using either the XlinkX or Sequest HT. For XlinkX searches, the database was searched for fully tryptic peptides, allowing for a maximum of 2 missed cleavages and a minimum peptide length of 5 amino acids. Searches were performed with a static modification of +57.021 Da on cysteine and a dynamic modification of +15.995 Da on methionine residues. Precursor mass tolerance, FTMS fragment mass tolerance, and ITMS Fragment tolerance, were set to 10 ppm, 20 ppm, and 0.5 Da, respectively. Xlink Validator FDR threshold was set to 0.01. For Sequest HT searches, the database was searched for fully tryptic peptides, allowing for a maximum of 2 missed cleavages and a minimum peptide length of 6 amino acids. Searches were performed with a static modification of +57.021 Da on cysteine, a dynamic modification of +15.995 Da on methionine residues, a dynamic modification of +176.014 Da on lysine residues (water-quenched DSSO monoadduct), and a dynamic modification of 279.078 Da on lysine residues (Tris-quenched DSSO monoadduct). Precursor mass tolerance and fragment mass tolerance were set to 10 ppm and 0.6 Da, respectively.

### Cloning ORFs into Plasmid Vectors

cDNA was prepared from human placental total RNA (Clontech) using the iScript cDNA synthesis kit (Bio-Rad). The coding sequences of MHAP1 (NP_001001436), MIER1b (NM_001077700.2), and MIER3-2 (NM_001297599.2) were amplified from cDNA using the PCR primers listed in Table S5 and cloned into either pFN21A or CMVd2 pcDNA5/FRT PacI PmeI as previously described ^17^. A codon-optimized ORF coding for SAP30 (NP_003855) was synthesized and subcloned into pFN21A PacI PmeI. To add a N-terminal HaloTag^®^, SIN3A was then excised with SgfI and PmeI, and cloned into SgfI and Eco53KI sites of pFC21A (Promega). SIN3A, with an in-frame N-terminal HaloTag^®^, was excised and cloned into MluI and NotI sites of pcDNA5/FRT Mammalian Expression Vector (Thermo Fisher). Halo-HDAC1 and HDAC1-Halo were constructed by amplifying with primers described in Table S5. The amplicon was digested with SgfI and PmeI and cloned into SgfI and Eco53KI sites of pFN21A or pFC14A. The CMVd2 promoter, HDAC1 ORF, and HaloTag^®^ in pFC16K were then amplified with primers described in Table S5. The amplicon was subsequently digested and cloned into MluI and NotI sites of pcDNA5/FRT mammalian expression vector. Halo-HDAC2 and HDAC2-Halo was created with a codon optimized sequence encoding HDAC2 synthesized commercially in the vector pUCIDT-AMP, excised by digestion with SgfI and PmeI, and then subcloned between the SgfI and PmeI sites of pFN21A or pFC14A. This sequence was amplified with primers described in Suppl. Table 9. The resulting amplicon was digested with SgfI and PmeI then cloned into SgfI and Eco53KI sites of pFC14A as previously described ^17^. The CMV promoter, HDAC1 ORF, and HaloTag^®^ in pFC14A were then amplified with primers described in Table S5. The amplicon was subsequently digested and cloned into MluI and NotI sites of pcDNA5/FRT mammalian expression vector.

### Western Blotting Analysis

The gel filtration samples were resolved on 10% SDS-PAGE gels. Gels were transferred to a PVDF membrane and blocked for one hour at 4 °C in 5% milk in Tris-buffered saline (TBS) and 0.1 % Tween-20. Primary antibodies were diluted in 5% milk in TBS and 0.1%Tween-20 and incubated overnight at 4°C. The following antibodies were used: Rabbit anti-HaloTag^®^ polyclonal (G9281) and anti-FLAG^®^ M2 mouse monoclonal primary antibodies; with IRDye^®^ 800CW labeled goat anti-Rabbit (926-3211) and IRDye^®^ 680LT labeled goat anti-Mouse (926-68020) secondary antibodies.

### Experimental Design and Statistical Rationale

For the MudPIT mass spectrometry analysis, DTASelect and Contrast ^63^ were used to filter results (minimum peptide length of 7 amino acid), and NSAFv7^64^ was used to calculate label-free quantitative dNSAF values as previously described ^17^. To identify high-confidence interaction partners, QSPEC v1.3.5^65^ was used to filter prey proteins. A high-confidence interaction was established when a protein was identified with log_2_FC >2, and FDR_up_ < 0.05 (or log_2_FC >3, Figure 2C) compared with controls expressing the Halo tag alone listed in Suppl. Table 5. The numbers of biological replicates used for each bait from transiently transfected cells were as follows (Suppl. Table S5): 7 x Halo-control; 4 x Halo-MHAP1; 4 x Halo-MIER1b; 4 x Halo-MIER3-2; 3 x HDAC1-Halo; 3 x HDAC2-Halo; 3 x Halo-SIN3A; 3 x Halo-SUDS3; 3 x Halo-SAP30. The mean spectral FDR for the 53 total MudPIT runs was 0.281% and the mean protein FDR was 0.766% as reported in Table S1. We have previously reported some of the MudPIT mass spectrometry data used in this study, as detailed in Table S5. Mass spectrometry data has also been deposited to the PeptideAtlas repository^66^ (www.peptideatlas.org), or in the MassIVE repository ^67^(http://massive.ucsd.edu); the identifiers for each dataset are listed in Table S5. For the XL-MS analysis, Xlink Validator FDR threshold was set to 0.01. Crosslinked peptides were identified from biological replicates for each bait as follows: 3 (HDAC1-Halo); 3 (HDAC2-Halo). A summary of the mass spectrometry runs used in this study is presented in Table S5.

### Fluorescence Microscopy

Imaging of Flp-In™-293 cell lines stably expressing HaloTag^®^ fusion proteins was performed as described previously ^17^. In brief, cells were seeded at 40% confluency in 35 mm MatTek glass bottom dishes (MatTek Corporation) containing DMEM supplemented with penicillin-streptomycin solution, GlutaMAX supplement, and FBS to a final concentration of 10%. Cells were first cultured for ∼24 hours at 37°C in 5% CO_2_. Affinity tagged proteins were fluorescently labeled with HaloTag^®^ TMR ligand (Promega), SNAP-Cell^®^ 505-Star (New England Biolabs), or both ligands according to the manufacturer’s instructions. Cells were stained with Hoechst 33258 solution (Sigma Aldrich) for 1 hour to label nuclei prior to imaging with a UltraVIEW VoX spinning disk confocal microscope (Perkin Elmer) with a 40x plan-apochromat (NA 1.2) objective. We excited TMR and Hoeschst with a 561 and 405 nm laser, respectively. All experiments implemented a multiband dichroic filter (405/488/561/640 nm) and a multiband emission filter (415-475/580-650 nm). Laser power was adjusted to maximize image quality, and brightness was adjusted using ImageJ/Fiji (http://fiji.sc) software as necessary.

Imaging experiments using transiently transfected 293T cells were performed essentially as described previously ^17^. 293T cells were plated to 40% confluency in MatTek 35 mm glass bottom culture dishes (MatTek Corporation) and cultured for 24 hours at 37°C in 5% CO_2_. Cells were then transfected or co-transfected with plasmids expressing the Halo-tagged, or SNAP-tagged protein of interest, as indicated. Affinity tagged proteins were fluorescently labeled with either HaloTag^®^ TMR™ Ligand (Promega) or SNAP-Cell^®^ 505-Star (New England Biolabs), or with both ligands according to the manufacturer’s instructions. Cells were stained with Hoechst 33258 solution (Sigma Aldrich) for 1 hour to label nuclei prior to imaging with an LSM-700 confocal microscope (Zeiss) and 40X plan-apochromat (NA 1.4) objective microscope. TMR was excited at 561 nm and collected through 578-696 nm, and Hoechst was excited at 405 nm and collected through 410-556 nm using MBS 405. Laser power was adjusted to maximize image quality and brightness was adjusted using ImageJ/Fiji (http://fiji.sc) software as necessary.

### Acceptor Photobleaching Fluorescence Resonance Energy Transfer

AP-FRET was performed essentially as described previously ^24,25^. Data was acquired on an Ultraview VoX spinning disc microscope (Perkin Elmer), equipped with a CSU-X1 spinning disc head (Yokogawa) recorded with an EMCCD camera (C1900-23B Hamamatsu) and a 100x (1.46 NA) alpha Plan-Apochromat objective (Zeiss) with Volocity software v.6.4.0. SNAP-Cell^®^ 505-Star (green channel) was excited with the 488 nm laser and emission collected with a 500–550 nm filter. HaloTag^®^ TMR (red channel) was excited with the 561 nm laser and emission collected with a 415–475 nm, 580– 650 nm dual band emission filter. Integration times for each channel were set as needed. We identified cells that contained both green and red fluorophores, collected 5 images in the green and red channels, bleached acceptor nuclei with 561nm laser at 100% power for 10 iterations, and then collected 5 more images after accepter photobleaching. This was repeated until data collection was complete (10–25 cells per sample). Using ImageJ/Fiji (http://fiji.sc) software to analyze the data, we first split the green and red channels and subtracted the background separately using the plug-in *roi average subtract jru v1*. Nuclei were outlined using the roi selection tool in ImageJ, and intensity measurements were taken using the plugin *create spectrum jru v1*. FRET efficiency was calculated by

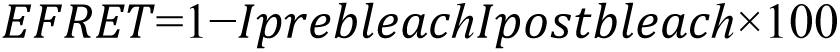

where I is the intensity from the donor channel averaged from the 5 images collected before and after photobleaching.

### Deacetylase Activity Assay

Deacetylase activity assays were performed as previously described ^17^. Briefly, approximately 1 x 10^7^ 293T cells were plated in 150 mm dishes and cultured in 25 mL DMEM with 10% fetal bovine serum and 1x GlutaMAX Supplement. Twenty-four hours after seeding, cells were transfected with 7.5 µg plasmid DNA, 7.5 µL Plus Reagent, and 50 µL Lipofectamine LTX diluted in 6.6 mL OptiMEM. Cells were harvested after an additional 48 h of culture. Two mg of whole cell extract were added to 100 µL of washed Magne^®^ HaloTag^®^ Beads slurry and incubated at 4°C for 2 h. Beads were washed 4 times with 1 mL cold TBS pH 7.4 + 0.05% Igepal CA-630 (Sigma Aldrich). Protein was eluted with 5 units AcTEV™ Protease (Thermo Fisher) in 100 µL of 50 mM Tris-HCl pH 8.0, 0.5 mM EDTA, 1 mM DTT for 16 h at 4 °C. Reactions containing 10 µl eluate were diluted with 0.1 mM Boc-Lys(Ac)-AMC, 32.5 µL TBS (25 mM Tris, 150 mM NaCl, 2mM KCl (pH 7.4) with or without 10 µM suberanilohydroxamic acid (SAHA) (Cayman Chemical) for 1 hour at 37°C. Reactions were quenched with 10 µM SAHA at 37°C for 5 min. Trypsin from porcine pancreas (Sigma Aldrich) was added to the reaction for a final concentration of 5 mg/mL for 1 hour at 37°C. Fluorescence was measured with a Tecan Infinite 200 PRO (excitation wavelength 355 nm, emission wavelength 460 nm). Calculations of deacetylase activity are reported in Table S2.

### Downstream Analysis of Affinity Purification Mass Spectrometry

Topological Scoring was performed for all bait proteins as described previously ^36^. Briefly, the average spectral counts for each bait from each AP-MS experiments were inputted into the TopS Shiny application (https://github.com/sardium/TopS). This application calculates the TopS scores, which indicate the likelihood of binding, by utilizing the average spectral counts of each bait across all baits. Hierarchical Clustering was performed on the QSPEC scores for each bait in which the log_2_(Fold Chage) was determined. This was completed in the R program heatmaply (https://talgalili.github.io/heatmaply/articles/heatmaply.html). Cytoscape (https://cytoscape.org/) was used to visualize relationships between significant interactions (significantly enriched with log2FC >2, FDRup <0.05) in at least one of the affinity purifications.

### Prediction of protein Structure Using AlphaFold

To predict the three-dimensional (3D) structure of HDAC2, MIER1, and MHAP1 (currently known as C16orf87 in databases) in various protein complexes, we used the AlphaFold3 protein structure prediction server ^10,11^. AlphaFold is a deep learning-based tool that provides highly accurate structural models by leveraging sequence-based input data. We first retrieved the amino acid sequence of each protein from the UniProt database (https://www.uniprot.org/). For the dimeric models, we input the sequence of HDAC2 along with the sequence of its interacting partner (MIER1 or MHAP1) or MIER1 and MHAP1 into AlphaFold to predict the corresponding structures. The AlphaFold server was run separately for HDAC2:MIER1, HDAC2:MHAP1, and MIER1:MHAP1, where each protein was modeled independently. To generate the HDAC2:MIER1:MHAP1 trimer, the sequences of HDAC2, MIER1, and MHAP1 were combined, and the AlphaFold Multimer ^10^ server was used to predict the structure of the entire trimeric complex. The resulting structural models were predicted using the default settings of the AlphaFold pipeline.

### Protein Structure Prediction and Docking

Following AlphaFold-only prediction, we modeled protein structures and intermolecular interactions using a *de novo* protein structure prediction and docking workflow as previously described in ^25,68^ and summarized in Fig. S3A. Initially, the amino acid sequence of the first isoform of a protein (from Uniprot) and, when available, PDB structural references were used as input for the initial prediction method, I-TASSER, ^20,21^. We used PDB template 3MAX for HDAC2 to guide the prediction with I-TASSER. Next, we assessed the predicted model for accuracy based on the distances measured for the intramolecular crosslinks, using a maximum of 37 Å between the a-C of the DSSO-crosslinked lysine residues (as described by ^25^). We measured a-C distances using both Euclidian distance (ED) in Chimera (https://www.cgl.ucsf.edu/chimera/) ^69^ and xWalk (https://www.xwalk.org/) ^70^, as well as Solvent Accessible Surface distance (SASD) in xWalk. We also calculated XL Scores for the initial model as described in ^68^. Briefly, this score counts the number of satisfied XLs for each model before the distance cut off. The models with the highest XL Scores were then moved on to Step III and inputted into I-TASSER along with their satisfied XLs as distance constraints. These XL-adjusted models were subjected to distance measurements between pairs of crosslinked lysine residues and XL scores were recalculated. A final output was then selected to be used for the molecular docking.

### Protein Docking

We performed protein docking in HADDOCK 2.2 (https://www.bonvinlab.org/software/haddock2.2/) ^22^. Prior to docking we used CPORT (https://alcazar.science.uu.nl/services/CPORT/) to determine and further guide the docking to potentially interacting residues, as described previously ^43^. Within HADDOCK 2.2, each satisfied XL was entered, “Center of Mass restraints” were turned on, CPORT analyses were for active and passive residues, and all other settings were selected for default. Our final model clusters were selected based on the best score measured by HADDOCK. If the clusters existed within a standard deviation of each other, we examined all clusters within that standard deviation. To assess the best model, we completed XL score analysis through measuring Chimera distances and then selected the best model from the best XL score.

### MHAP1 Protein Model Selection

MHAP1 (currently known as C16orf87 in databases) had no intra-protein XLs from which we could assess, guide, and refine a *de novo* structure prediction. We then selected the models with the best scores as calculated by I-TASSER ^20,21^. We next performed HDAC1:MIER1:MHAP docking with the best I-TASSER MHAP1 model and all intermolecular crosslinks between three proteins we detected in the HDAC2, MIER1, and MHAP1 XL-MS analyses. The distances between intermolecular crosslinks on the predicted complex were measured as described above.

### Structure Analysis Using Customized Python Scripts

The predicted 3D structures were processed and analyzed using custom Python scripts. Structural comparisons were conducted by superimposing the predicted HDAC2 models from different complexes, focusing on the IDR in the C-terminal and globular middle region of the protein. This allowed the identification of structural shifts or flexibility differences in the multiple dimeric complexes formed by HDAC2, MIER1, and MHAP1. The key steps in the structure analysis involved lysine-lysine interactions that were predicted by identifying pairs of lysine residues in proximity within 30 Å of each other. In these interactions, we calculated the distance between lysine residues and generated interaction maps for each complex. Residue contact maps were also generated by calculating the Euclidian distances of each pair of interacting residues within 30 Å of each other. Contact maps were plotted to compare the density of residue-residue interactions in different complexes, including the dimers and the trimer. The maps were then used to assess the extent of structural differences between the complexes. Finally, all 3D structures were visualized using ChimeraX ^69^ and VMD ^71^ packages to facilitate direct comparisons of multiple conformations in different complexes. The structures were superimposed to isolate the shifts and the structural flexibility from different complexes. The superimpositions allowed us to evaluate structural variations in the conserved regions of individual protein upon forming a complex and to compare the results with the ones obtained using an integrative approach.

## Acknowledgments

Research reported in this publication was supported by the National Institute of General Medical Sciences of the National Institutes of Health under award numbers F31GM131536 (C.G.K.), T32GM138077 (J.C.), R35GM118068 (J.L.W.), and R35GM145240 (M.P.W.) and the Stowers Institute for Medical Research. The content is solely the responsibility of the authors and does not necessarily represent the official views of the National Institutes of Health. This work was performed as part of C.G.K.’s research thesis at the Graduate School of the Stowers Institute for Medical Research.

## Author Contributions

Conceptualization: JN, CGK, MPW

Investigation: JN, CGK, JC, RZ, YZ

Formal Analysis: JN, CGK, RZ, YZ, LF, MPW

Resources: JN, CGK, YZ, CASB, LF, MPW

Software: JN, CGK, JC

Data Curation: JN, CGK, JC, LF

Writing: JN, CGK, LF, MPW

Supervision: LF, JLW, MPW

Funding acquisition: CGK, JLW, MPW

## Data and Materials availability

Mass spectrometry data may be accessed through Proteome Xchange via the MassIVE repository using the accession numbers reported in Supp. Table S5. After publication, original mass spectrometry data underlying this manuscript may also be accessed from the Stowers Original Data Repository at https://www.stowers.org/research/publications/LIBPB-1566.

---

**Supplemental Figure S1.**
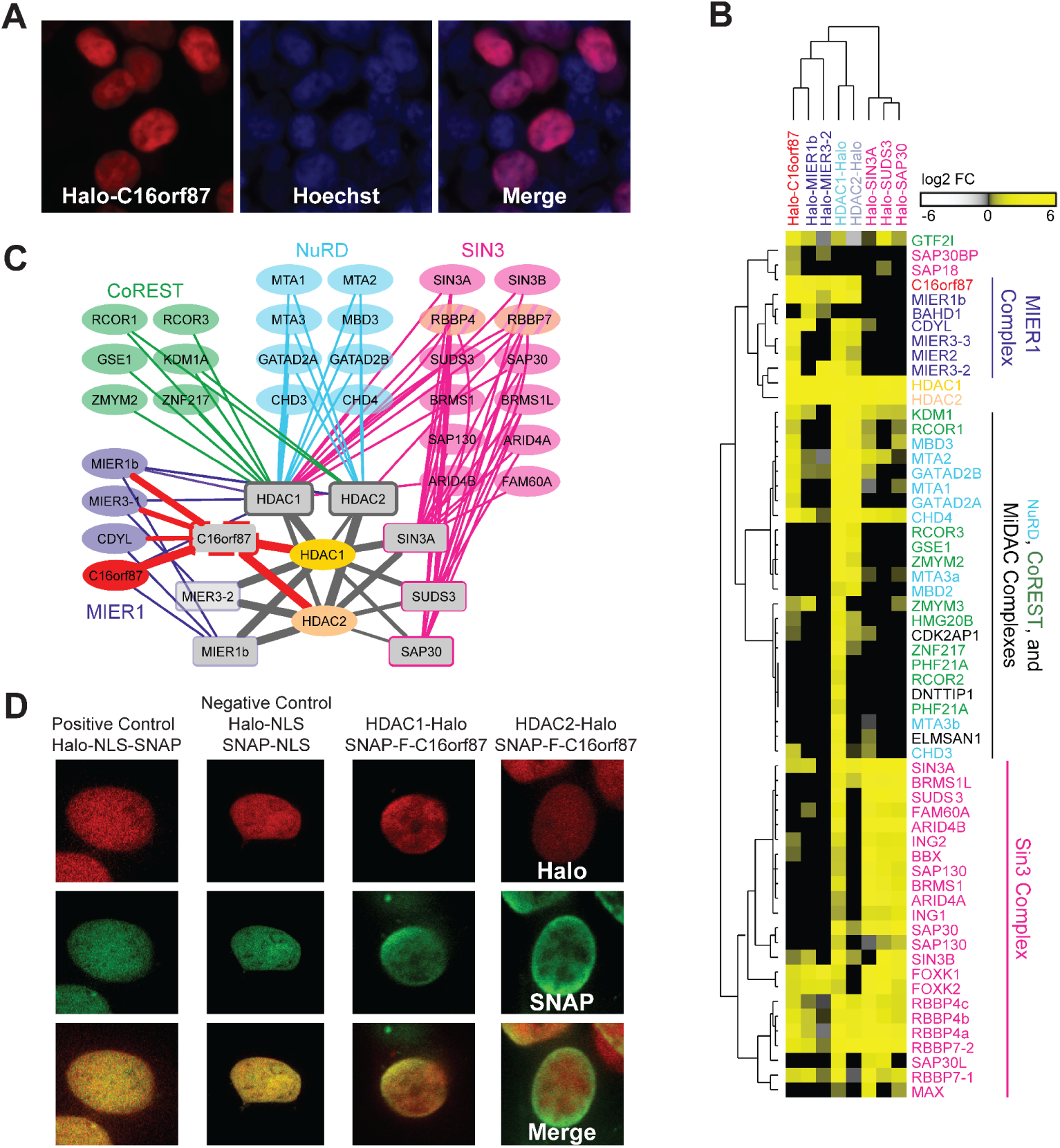
C16orf87 localization and interaction network. **A**. Nuclear Localization of HALO-C16orf87 in HEK293T. Halo-tagged proteins are labeled with HaloTag® TMR ligand (red). Nuclei are stained with Hoechst dye (blue). **B**. Heatmap plotting relative protein abundance expressed as log2FC with the hierarchical bi-clustering of the 8 Halo-tagged baits: C16orf87, MIER1b, MIER1-2, HDAC1, HDAC2, SIN3A, SUDS3, SAP30, with the brightest yellow indicating high log2FC (Suppl. Table S1A). **C**. A network analysis was performed using Cytoscape on members of HDAC1/2 associated complexes significantly enriched with at least one of the bait proteins compared to controls (log2FC > 3, FDRup< 0.05; Suppl. Table S1A). The Halo-tagged bait proteins (C16orf87, MIER1b, MIER1-2, HDAC1, HDAC2, SIN3A, SUDS3, and SAP30) are source nodes in gray, with SIN3A associated proteins in pink, NuRD associated proteins in light blue, CoREST associated proteins in green, MIER associated proteins in purple, HDAC1/2 (in light yellow/orange, and C16orf87 in red. **D**. Representative images of acceptor and donor fluorescence for each sample in the AP-FRET experiments. SNAP-F-C16orf87 is labeled with the SNAP-Cell® 505-Star ligand, while HDAC1/2-Halo are labeled with the HaloTag® MR ligand (see data in Suppl. Table S2A).

**Supplemental Figure S2.**
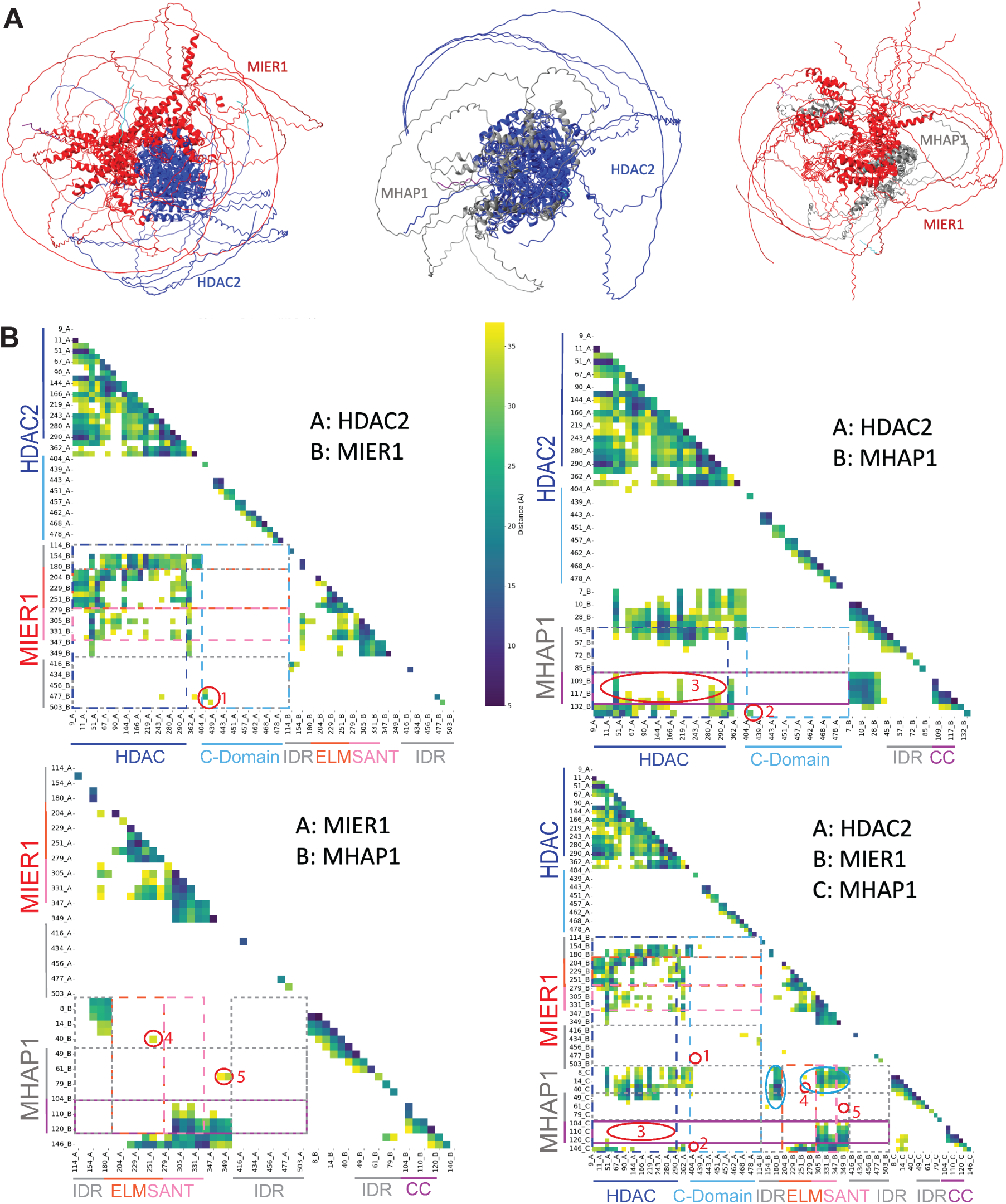
Structural Comparison of Alpha-Fold-Based Dimer and Trimer Models. **A.** Five AF-predicted assemblies are superimposed for each of the dimeric models. **B.** Lysine-lysine interactions within 30Å are plotted for each model. Known structural or IDR domains are indicated by colored lines along the axes. Numbered areas of difference between the dimers and trimer are circled in red and blue.

**Supplemental Figure S3.**
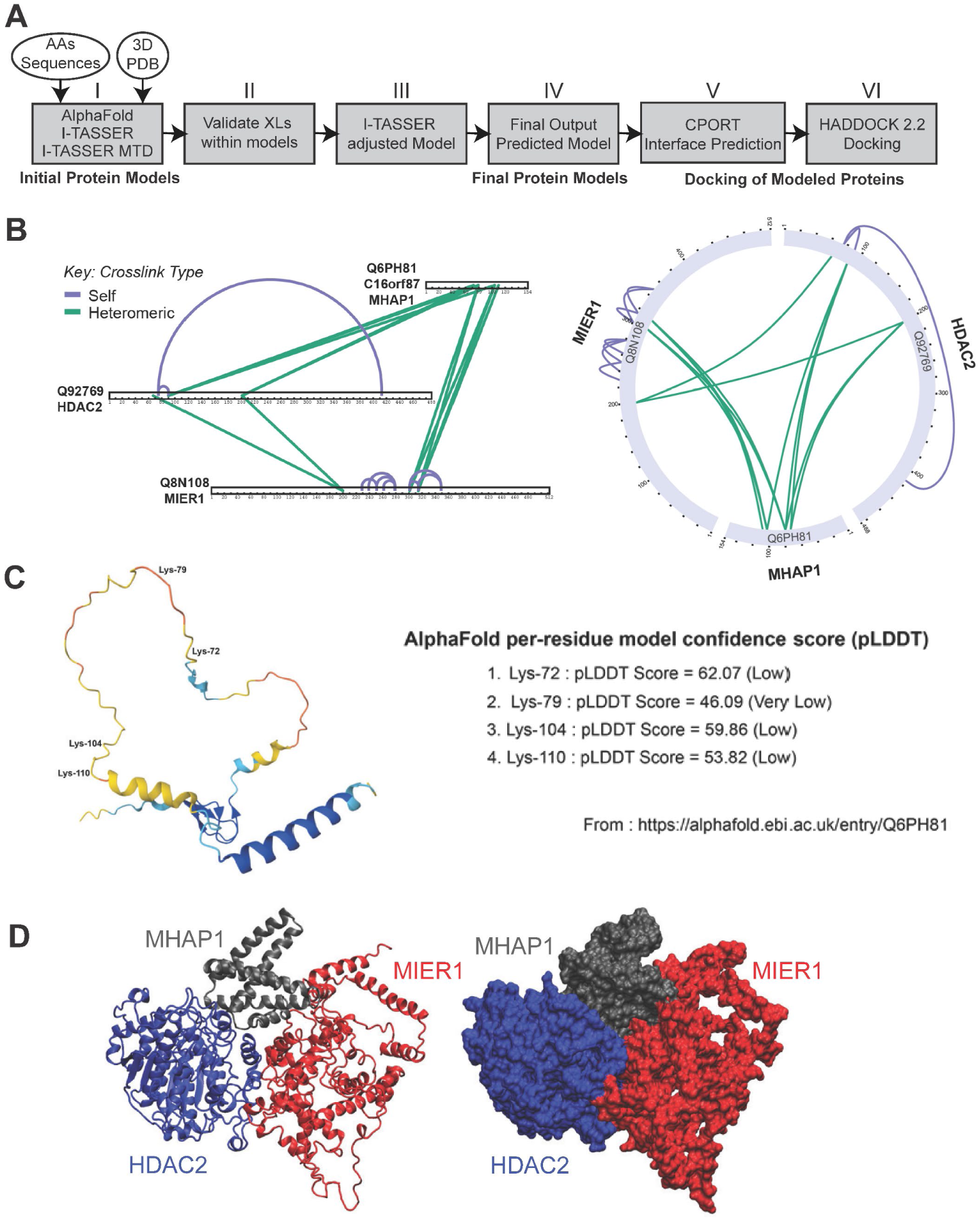
XLMS-Guided Integrative Structural Modeling of the MHAP1:MIER1:HDAC2 Trimer. **A.** *de novo* protein prediction and docking workflow implemented to model the 3D structures of individual proteins and their interaction interfaces and docking in the final complex. **B.** Linear and Circos plots of the crosslinks observed in the XLMS analysis of MHAP1-Halo purifications (see Supp. Table S3). **C.** The four lysine residues found crosslinked in UPF0547 protein C16orf87 in this study are listed with their AlphaFold per-residues scores (from AF-Q6PH81-F1-v4). **D.** Trimeric assembly predicted after step VI of the ISM workflow for HDAC1, MIER1, and MHAP1 (visualized in Chimera as ribbon and space-filled representations).

**Supplemental Figure S4.**
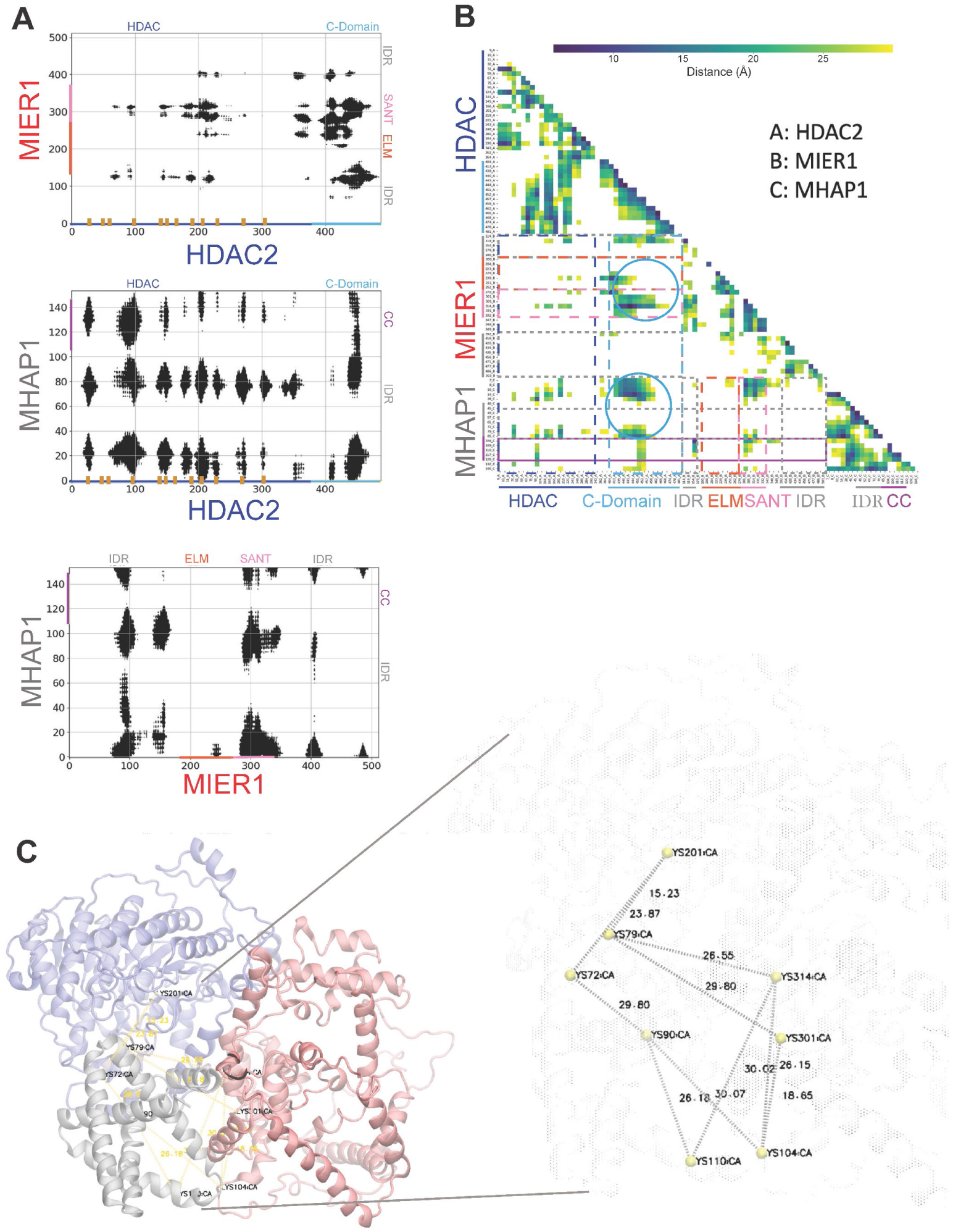
Contact Maps and Core Distances within the ISM-Trimer. **A-B.** Binary residue contact maps (with HDAC ligand binding sites shown as brown tick marks along the x-axis) and heatmap of lysine-lysine distances (within 30 Å) with key interacting hotspots between HDAC C-domain and MIER1 and MHAP1 circled in blue. **C.** Euclidian distances between the α-carbons of the lysine residues (circled in **B**.) forming the stable core of the ISM-predicted ternary assembly, with a scaled-up area of the distances measurements shown on the right.

**Supplemental Figure S5.**
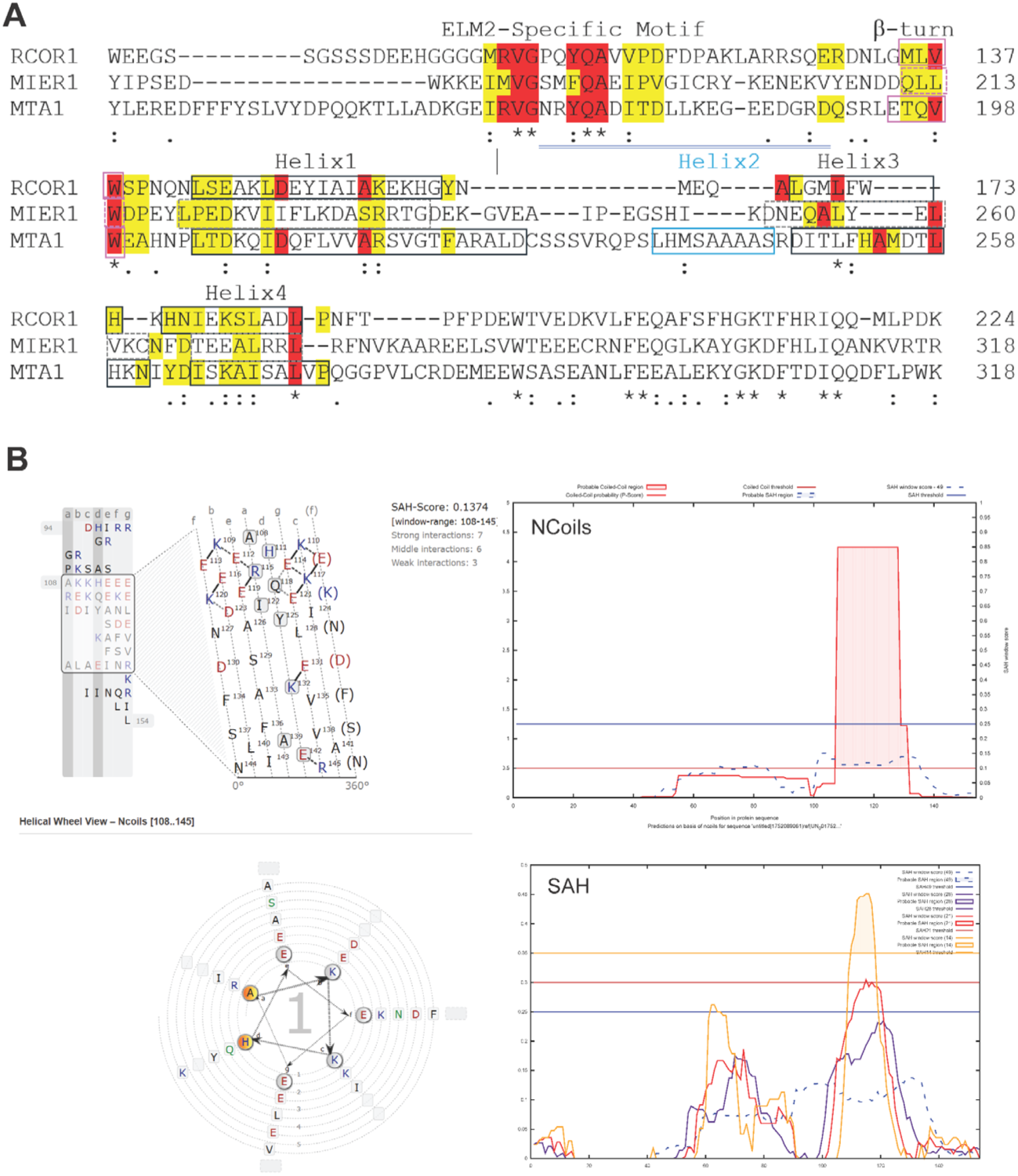
Secondary Structures Analyses of MIER1 ELM2 and MHAP1 CC Domains. **A.** Clustal Omega sequence alignment of ELM2 domains in the RCOR1–3, MIER1, and MTA1– 3 proteins. Identical residues are shown in red and conserved residues are shown in yellow. The predicted secondary structures of RCOR1 and the secondary structures of MTA1 are indicated as boxed amino acids, with helix numbers corresponding to the solved MTA1 ELM2 domain. The predicted secondary structures for MIER1 are dashed. **B.** N-Coils helical net and wheel views and prediction results from the MHAP1 sequence to distinguish coiled-coil (CC) from stable a-helix (SAH) domains, all obtained from the Waggawagga prediction algorithm (https://waggawagga.motorprotein.de/).

